# The longer the better? General skill but not probabilistic learning improves with the duration of short rest periods

**DOI:** 10.1101/2020.05.12.090886

**Authors:** Lison Fanuel, Claire Plèche, Teodóra Vékony, Romain Quentin, Karolina Janacsek, Dezso Nemeth

**Affiliations:** Lyon Neuroscience Research Center (CRNL), INSERM, CNRS, Université Claude Bernard Lyon 1, Lyon, France; Department of Neurology, University of Szeged, Szeged, Hungary; Human Cortical Physiology and Neurorehabilitation Section, NINDS, NIH, Bethesda, Maryland, USA; School of Human Sciences, Faculty of Education, Health and Human Sciences, University of Greenwich, Old Royal Naval College, Park Row, 150 Dreadnought, SE10 9LS, London, United Kingdom; Brain, Memory and Language Research Group, Institute of Cognitive Neuroscience and Psychology, Research Centre for Natural Sciences, Magyar tudósok körútja 2., H–1117, Budapest, Hungary; Institute of Psychology, ELTE Eötvös Loránd University, Izabella utca 46., H-1064, Budapest, Hungary

**Keywords:** consolidation, implicit learning, probabilistic learning, offline learning

## Abstract

Memory consolidation has mainly been investigated for extended periods, from hours to days. Recent studies suggest that memory consolidation can also occur within shorter periods, from minutes to seconds. Our study aimed at determining (1) whether short rest periods lead to improvements in implicit probabilistic sequence learning and (2) whether length of rest duration influences such offline improvements. Participants performed an implicit probabilistic sequence learning task throughout 45 blocks. Between blocks, participants were allowed to rest and then to continue the task in their pace. The overall reaction times (general skill learning) shortened from pre- to post-rest periods, and this improvement was increased for longer rest durations. However, probabilistic sequences knowledge decreased in these periods, and this decrement was not related to the length of rest duration. These results suggest that (1) general skill learning but not probabilistic sequence knowledge benefits from short rest periods and, possibly, from memory consolidation, (2) ultra-fast offline improvements in general skills, but not forgetting in probabilistic sequence knowledge, are time-dependent. Overall, our findings highlight that ultra-fast consolidation differently affects distinct cognitive processes.

## Introduction

Taking a break during a learning period may facilitate the acquisition of new perceptual and motor skills (e.g., perceptual discrimination or finger tapping) and also benefit more complex cognitive skills, such as solving mathematical problems (e.g., Fischer et al., 2002; Stickgold et al., 2000; Stickgold & Walker, 2004; Walker et al., 2002). During rest periods (i.e., between two learning sessions), our brain strengthens memories through consolidation, potentially leading to performance improvements (e.g., Robertson, Pascual-Leone, & Miall, 2004). So far, consolidation processes have mainly been investigated on extended periods following learning, such as days or hours (Squire et al., 2015 for a review). Recent studies showed that shorter rest periods, within a single learning session, also benefit performance (Bönstrup et al., 2019; Du et al., 2016) and was referred to as ultra-fast offline improvement (Robertson, 2019). These studies focused on the acquisition of new motor skills. In the present study, we wondered whether short rest periods could more broadly benefit the development of new cognitive or social abilities. As implicit probabilistic sequence learning underlies the acquisition of motor, cognitive and social skills (Lieberman, 2000; Nemeth et al., 2011; Romano Bergstrom et al., 2012; Ullman, 2016), the present study investigated whether and how this type of learning also benefits from short rest periods.

The first empirical evidence for ultra-fast offline improvements was provided for motor sequence learning of deterministic sequences during ten-second rest periods (Bönstrup et al., 2019). In that study, participants learned a finger-tapping sequence, alternating between ten seconds of practice and ten seconds of rest. Performance improvements over practice and rest periods were separately measured. Increases in performance over rest periods considerably contributed to the overall learning of the tapping task, suggesting the strengthening of just-practiced skills during rest periods. Concomitant magnetoencephalographic measures further highlighted modulation in beta-band frequency during rest periods. Beta-band oscillations are associated with reactivation of previous practice-related activity (Maquet et al., 2000; Ramanathan et al., 2015; see also Spitzer & Haegens, 2017 for a review), also referred to as memory replay (Cohen et al., 2015). Bönstrup et al. (2019)’s results thus suggest that ultra-fast offline improvements can be related to the reactivation of memory traces (see also Robertson, 2019 for a similar hypothesis).

To date, investigation of ultra-fast offline improvements has focused on explicit motor sequence learning, suggesting that short rest periods benefit acquisition of new motor skills. As consolidation processes seem to vary depending on the awareness of learning (Robertson, Pascual-Leone, & Press, 2004), we wondered whether ultra-fast offline improvements could extend to implicit probabilistic sequence learning. Implicit probabilistic sequence learning can be described as the development of knowledge about regularities embedded in the environment without awareness nor intention of learning (e.g., Cleeremans & Jiménez, 1998; Howard et al., 2004). This sort of learning is involved in acquisition of new motor, cognitive and social skills (Lieberman, 2000; Nemeth et al., 2011; Romano Bergstrom et al., 2012; Ullman, 2016). Ultra-fast offline improvements in implicit probabilistic sequence learning would suggest that short rest periods also benefit the acquisition of cognitive and social abilities.

In the present study, we wondered whether short periods of rest could lead to ultra-fast offline improvements in implicit acquisition of probabilistic sequence knowledge. To address this question, we used the Alternating Serial Reaction Time (ASRT) task (e.g., Howard et al., 2004; Song et al., 2007). In this paradigm, an array of four positions were presented on the screen, and each position was mapped to a specific response key. On each trial, one of the positions was filled, and the participant had to press the corresponding key as fast and accurately as they could. Importantly, without the participant’s awareness, the sequence of events followed a predictable pattern that was embedded in noise (i.e., presented among random positions). Participants were offered to rest after each block (corresponding to 85 trials) and resumed the task whenever ready. Our experimental design with self-paced rest periods overcame the limitations of the previous studies using fixed periods by allowing us to directly measured how length of rest periods affected learning performance. Probabilistic sequence knowledge was evaluated by comparing the speed and accuracy of responses depending on the items’ probability of occurrence (high-probability or low-probability) was measured before and after each rest period. An increase in probabilistic sequence knowledge over the rest period would reflect an ultra-fast offline improvement of implicit probabilistic sequence. Besides inducing implicit probabilistic sequence learning, ASRT task allows to distinguish it from more general motor or visuomotor skill (further referred to as general skill) learning (Hallgató et al., 2013; Nemeth et al., 2010; Song et al., 2007). Based on previous findings of ultra-fast offline improvements in motor sequence learning (Bönstrup et al., 2019), we expected an increase in general skills (i.e., response speed and accuracy regardless of the probability of occurrence of items) over the rest period. Furthermore, we investigated whether rest period duration could influence ultra-fast offline improvements. If ultra-fast offline improvements occur over rest periods (either in implicit probabilistic sequence knowledge or in general skills), then longer rest periods would lead to greater offline improvement. On the contrary, if memory decay occurs during rest periods, we expected rest periods to impact the amount of memory decay.

## Method

### Participants

One hundred and eighty healthy young adults participated in this study (M_age_ = 21.64 years, SD_age_ = 4.11, M_education_ = 14.69 years, SD_education_ = 2.16, 152 females). All participants had normal or corrected-to-normal vision, and none of them reported a history of any neurological and/or psychiatric condition. Participants provided informed consent to the procedure before enrollment as approved by the institutional review board of the local research ethics committee. The study was approved by the United Ethical Review Committee for Research in Psychology (EPKEB) in Hungary (Approval number: 30/2012) and by the research ethics committee of Eötvös Loránd University, Budapest, Hungary. The study was conducted in accordance with the Declaration of Helsinki. Participants received course credits for taking part in the experiment. The dataset was previously used in Kóbor et al. (2017) and Török et al. (2017). These two articles explore different questions than the one reported here. Results constituting the present paper were not tested nor reported before.

### Alternating Serial Reaction Time Task

The Alternating Serial Reaction Time (ASRT) task was used to induce implicit probabilistic sequence learning (Howard et al., 2004; Song et al., 2007). Four empty circles were horizontally arranged on the screen. A stimulus (a drawing of a dog’s head) appeared in one of four circles (Figure 1.A.) (Nemeth et al., 2013). Participants were instructed to press the corresponding key (Z, C, B, or M on a QWERTY keyboard) as quickly and accurately as possible after the appearance of the stimulus. Participants used their left and right middle and index fingers to respond to the targets. The serial order of the four possible positions (coded as 1, 2, 3, and 4) in which target stimuli could appear was determined by an eight-element sequence. In this sequence, every second element appeared in the same order as the task progressed, while the other elements’ position was randomly chosen (e.g., 2 – *r* – 1 – *r* – 3 – *r* – 4 – *r*; where numbers refer to a predetermined location in one of the four locations and *r’*s refer to randomly chosen locations out of the four possible, Figure 1.B.). Six different sequences of predetermined elements were created and assigned to each subject in a permutated order.

**Figure 1.**
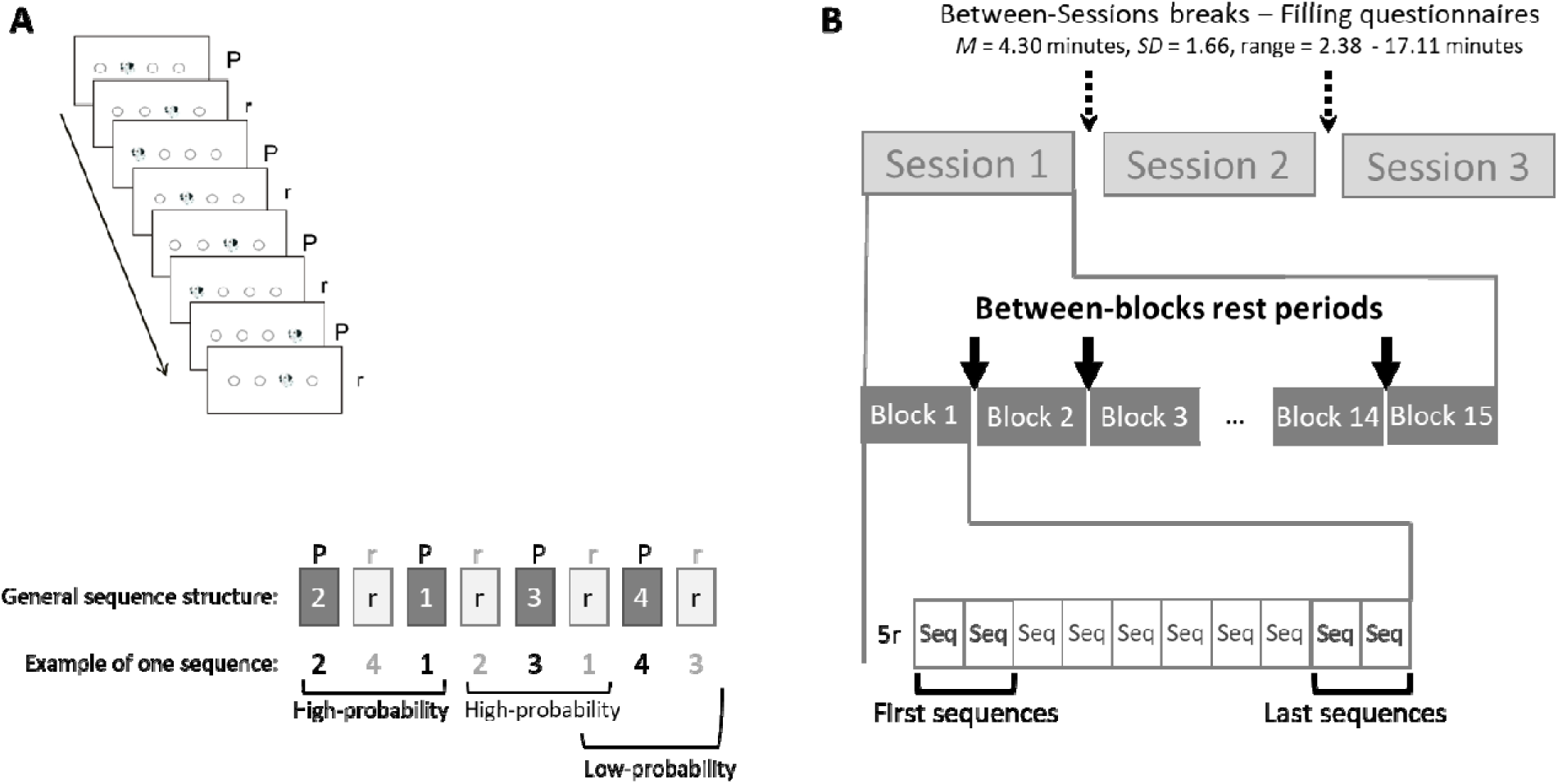
Schematic representation of (A) an ASRT sequence and (B) the overall structure of the task. Each sequence was composed of eight elements alternating between predetermined (P) and random (r). The experiment was divided into three sessions, each composed of 15 blocks. A rest period was offered after each block (arrows). Between-sessions breaks (dotted arrows) were discarded from analyses because participants filled questionnaires during this time. Only self-paced between-blocks rest periods (bold arrows) were included in the analyses. Each block was composed of five warm-up random trials (5r), followed by ten eight-element sequences (Seq). Brackets flag the two first and the two last sequences from which offline improvement scores were computed.

Due to the alternating sequence structure, some patterns of three consecutive elements (henceforth referred to as triplets) occurred with a greater probability than other ones (Figure 1.B). *Each element* was categorized as either the third element of a high- or a low-probability triplet. High-probability triplets could be either formed by predetermined elements or random ones. For instance, the probability that 4 – r – 2 occurred was of 62.5% (i.e., if the item 4 was the first triplet element, the item 2 had 50 % probability of occurring because it was a predetermined element plus 12.5% of chances to occur as a random element). The third element of less probable triplets (e.g., 1 – r – 2 and 4 – r – 3) could have only been random and was thus less predictable (e.g., if the first triplet element was the item 4, the item 3 had 12.5% of chances to occur). Low-probability triplets forming repetitions (e.g., 222) or trills (e.g., 232) were discarded from analyses as participants often show preexisting response tendencies to them. By eliminating these triplets, we could ascertain that any high- versus low-probability differences were due to learning and not to preexisting tendencies.

As the task progresses, participants usually become faster and more accurate for the high-probability triplets compared to the low-probability ones. Therefore, the task allows us to separate pure *implicit probabilistic sequence learning* (i.e., the difference between high- and low-probability triplets) from *general skill learning* (Song et al., 2007). General skill learning refers to changes in accuracy and response times independently from the probability of occurrence of the events (Hallgató et al., 2013).

### Procedure

The ASRT task was administered in three sessions, each containing 15 blocks (45 blocks in total). Each block consisted of 85 trials, corresponding to five warm-ups, random trials followed by the eight-element sequence repeated ten times (Figure 1.C.). Accuracy and response time (RT) were recorded for each element. Between each block, a rest was proposed, and participants resumed the task whenever they were ready. Between sessions, participants filled questionnaires. Thus, only between-block rest periods were included in the following analyses.

### Quantification and statistical analyses

To assess the impact of rest duration on probabilistic sequence learning, we measured the length of between-blocks rest periods as well as various indices of learning. We measured probabilistic sequence knowledge acquired across the whole experiment as well as at the beginning and the end of each block. We further provided a measure of offline gain in probabilistic sequence knowledge during each rest period.

### Between-blocks rest periods measure

The amount of time elapsed between the last response of block N and the key-press that started block N+1 was computed for each between-block rest period (M = 18.35 seconds, SD = 9.40 seconds, range = 15.39 to 480 seconds). This procedure resulted in 42 measures of between-blocks rest durations for each participant. Between-blocks rest durations were averaged for each participant (referred to as mean between-blocks rest periods). To account for possibly erroneous procedures (e.g., participant had to leave the room), participants whose average between-blocks rest durations that was below or above the conventional exclusion threshold of 2 SD above the mean were considered removed from the sample. Therefore, the following statistical analyses included 172 participants aged between 17 and 48 years (M_age_ = 21.63 years, SD_age_ = 4.16, M_education_ = 14.68 years, SD_education_ = 2.14, 146 females).

### General skills

General visuomotor skills were considered as the speed and accuracy of responses irrespective of the items’ probability. Thus, RT and accuracy measures independent from triplets’ probability were considered as an index of general skills. General skills index was calculated by computing mean accuracy and median RT for correct responses for the two first sequences and the two last sequences of each block, including all items of that sequences irrespective of the items’ probability. This procedure resulted in four scores for each block: median RT and mean accuracy for the first two sequences and median RT and mean accuracy for the last two sequences. Then, RT and accuracy scores were separately averaged for the first and last sequences for each participant.

### Probabilistic sequence knowledge

Probabilistic sequence knowledge was considered as the difference in performance depending on triplets’ probability of occurrence. To compute an index of probabilistic sequence knowledge, we first calculated mean accuracy and median RT for low- and high-probability triplets separately. Probabilistic sequence knowledge score consisted of the difference between the score for high-probability triplets minus the score for low-probability triplets. For RT, a lower score indicated larger probabilistic sequence knowledge for accuracy. For accuracy, the opposite was true: the greater score, the larger probabilistic sequence knowledge. Following this procedure, we measured (1) the *probabilistic sequence learning* based on RT and accuracy scores of the entire experiment and (2) the *probabilistic sequence knowledge* acquired at the beginning and the end of each block (akin index of general skills described previously). To do so, we first measured mean accuracy and median RT for correct responses for the two first sequences (i.e., first 14 trials after the five warm-up trials) and the two last sequences (i.e., last 16 trials) of each block and each triplet probability. This method resulted in eight scores for each block: median RT and mean accuracy for the first two sequences for low-probability items, for the first two sequences for high-probability triplets, and for the last two sequences for low-probability items, and the last two sequences for high-probability items. Four scores of probabilistic sequence knowledge were computed for each block: accuracy and RT indices for the first two blocks and the last two blocks. Then, probabilistic sequence knowledge for both RT and accuracy was separately averaged for the first and last sequences of the block and this for each participant.

### Offline modulations

Offline modulations were considered as a change (i.e., either an increase or a decrease) in RT or in accuracy between the last two sequences of a given block and the first two sequences of the next one (after warm-up trials). *Offline modulations in general skills* consisted of the difference in RT or accuracy scores between the first sequences of a block and the last sequences of the previous block irrespectively of triplet probability. *Offline modulations in probabilistic sequence knowledge* consisted of the difference of RT or accuracy indices of probabilistic knowledge (i.e., the difference between high- and low-probability triplets) between the first sequence of a block and the last sequence of the previous block. For RT indices, a negative difference shows a speeding up of responses and suggests offline improvement, while a positive difference shows a slowing down of responses and suggests memory decay. For accuracy, the opposite pattern is true. The calculation of offline modulations for both general skills and probabilistic sequence knowledge resulted in 42 offline scores that were averaged for each participant.

### Post-hoc subgrouping

To include between-blocks rest duration as a factor in the following analyses, we transformed the mean between-blocks rest duration into a categorical measure. Participants were divided into three groups around the 33^rd^ and 66^th^ percentile (16.91 seconds and 17.86 seconds): participants who had the shortest between-blocks rest periods (i.e., below 33^th^ percentile, *N* = 57, *M* = 16.52 seconds, *SD* = 0.22), participants who had the longest between-blocks rest period (i.e., above 66^th^ percentile, *N* = 59, *M* = 20.34 seconds, *SD* = 2.52), and participants in the median group (between 33^th^ percentile and 66^th^ percentile, *N* = 55, *M* = 17.34 seconds, *SD* = 0.27). Analyses using post-hoc subgrouping included participants who had the shortest and the longest between-blocks rest period and not participants from the median group to create a stronger separation between groups. No between-group differences emerged in demographic variables (Age, Education level, and Sex; see Supplementary material).

### Linear relationship between offline modulations and rest duration

We assessed the relationship between the ultra-fast offline modulations in general skills and probabilistic sequence knowledge and the duration of the between-blocks rest periods. For *between-participants analysis*, mean offline modulations in general skills and probabilistic sequence knowledge were calculated for both RT and accuracy for each participant. We tested their correlation with mean rest duration using frequentist Pearson’s and Bayesian correlations. To account for the inter-individual variability, we conducted *within-participant analyses*. Beforehand, aberrant data points were removed: between-block rest durations that were 2 SD above the participant’s mean rest duration were excluded from the sample. We removed 2 ± 0.81 between-block rest duration (range: 0 – 4) for each participant (i.e., 4.76% of the total amount of data points). For both RT and accuracy measures, we computed Pearson’s correlations between between-blocks rest duration and offline modulation (in general skills and probabilistic sequence knowledge) separately for each participant. The resulting correlation coefficients (Pearson’s r) were considered as an individual measure of the relationship between between-blocks rest duration and offline modulation in general skills and probabilistic sequence knowledge for RT and accuracy. Frequentists and Bayesian one-sample t-tests contrasting correlation coefficients to zero were conducted separately for each measure (i.e., RT measure for general skills, accuracy measure for general skills, RT measure for probabilistic sequence knowledge, accuracy measure for probabilistic sequence knowledge).

### Bayesian statistical analyses and guidelines for interpretation

In addition to classical frequentist statistics, Bayesian statistics were computed. Bayes factors (BF) were calculated for each model potentially fitting the data, that is for all main effects and combinations of factors (additive or interactive) included in the analysis. The BF_10_ associated with a given effect resulted from comparing all the models including the effect to all the models *not* including it (Etz & Wagenmakers, 2017). Thus, it reflects the probability of the inclusion of this effect averaged across all candidate models. When applicable, we reported the Bayes factor associated with an effect (BF_10_) as well as the most probable model (i.e., the model that fits best the data among all the possible models) and its associated Bayes factor (BF_M_). Importantly, a Bayes factor can give evidence towards the alternative hypothesis (H1) or the null hypothesis (H0). BF_10_ between 3 and 10 and above 10 is considered as moderate support and strong support for the alternative hypothesis, respectively (Lee & Wagenmakers, 2014). BF_10_ values between 1/3 and 1/10 and below 1/10 are considered as moderate support and strong support for the null hypothesis, respectively. BF_10_ values between 1/3 and 3 are regarded as ambiguous information (Etz et al., 2017; Lee & Wagenmakers, 2014; Wagenmakers, 2007). All statistical analyses were performed using JASP 0.11.1 (JASP Team, 2019) with the default settings.

## Results

### Did between-block rest periods influence general skills and probabilistic sequence knowledge?

To test whether ultra-fast offline improvements in general skill and implicit probabilistic sequence knowledge occurred during between-block rest periods and to evaluate the effect of between-block rest duration, mixed-design repeated-measures ANOVAs were run with Block (Last sequences of block n vs. First sequences of block n+1) as a within-participants factor and Group (Short vs. Long between-block rest duration, see Post-hoc subgrouping procedure in the method section) as a between-participants factor. Frequentists and Bayesian ANOVAs were conducted on RT and accuracy measures for both general skills and probabilistic sequence measures (Figure 2).

**Figure 2.**
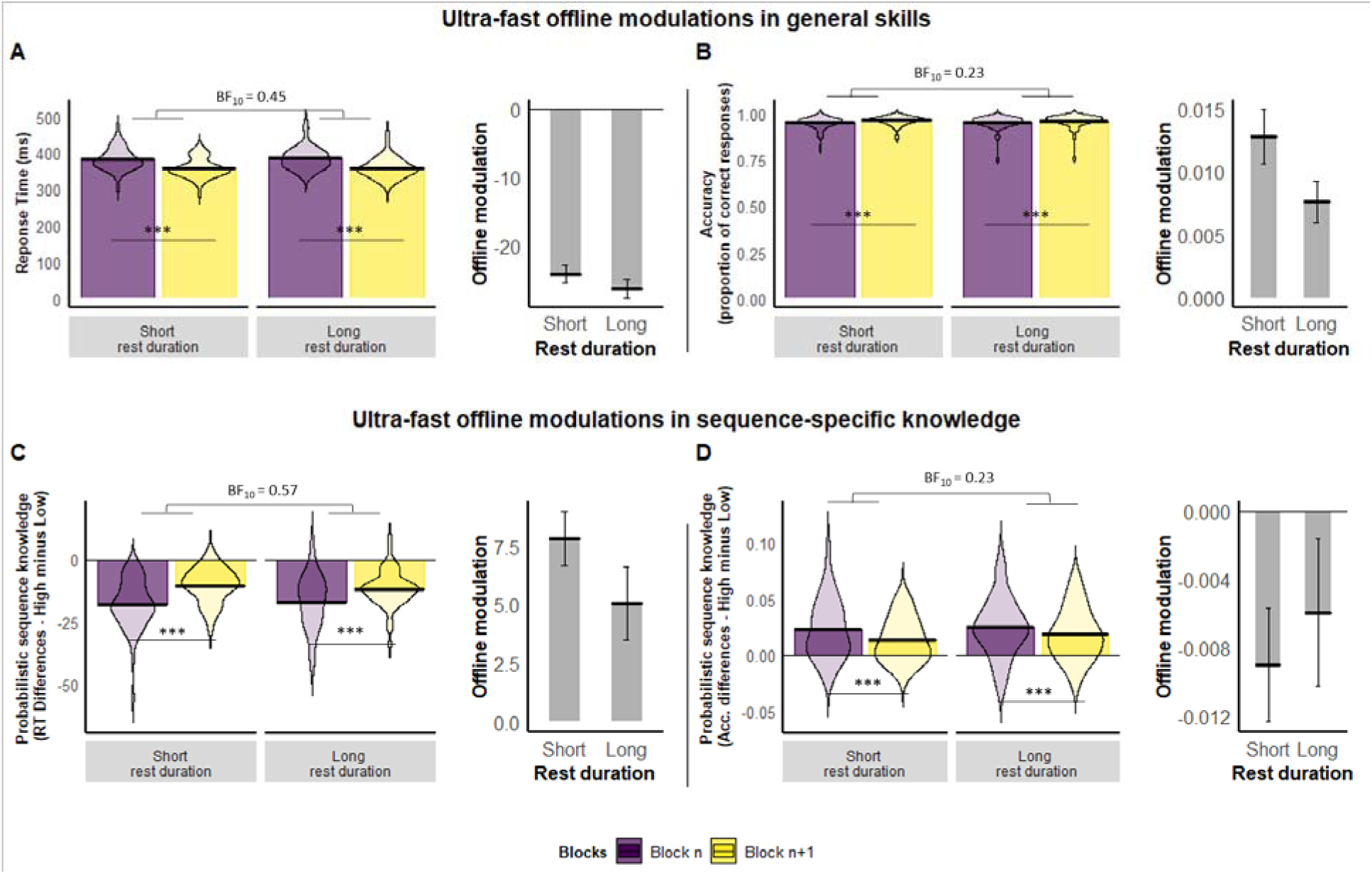
Offline modulations in general skills (A, B) and in probabilistic sequence knowledge (C, D). In each sub-figure, the left-sided plot corresponds to median response times in milliseconds (A), mean accuracy in the proportion of correct responses (B), or RT and accuracy measures of probabilistic sequence knowledge (C and D, respectively) as a function of Block (Block n: Last sequences of block n, Block n+1: First sequences of block n+1) and Between-blocks rest duration (Short vs. Long). The right-sided plot corresponds to offline modulation measures, that is the difference in knowledge indices (raw RT, raw accuracy, probabilistic sequence knowledge) between the first sequences of block n+1 and the last sequences of block n. For RT measures, negative modulation (i.e., speeding of responses) corresponds to offline improvement and a positive modulation (i.e., slowing of responses) corresponds to offline decrement. The opposite is true for accuracy measures. Response times were faster and accuracy higher in the first sequences of block n+1 than in the last sequences of block n, suggesting offline gains in general skills. However, probabilistic sequence knowledge decreased over between-block rest periods, suggesting an offline decay. Moreover, between-blocks rest duration did not influence neither offline gains in probabilistic sequence knowledge nor in general skill. Violin plots represent data distribution; black horizontal lines represent the mean across participants. Vertical error bars represent standard error. *** stands for p < .001. Bayes factor is reported for non-significant effects. BF < ⅓ shows evidence for the null hypothesis (see method section for more details).

### Ultra-fast offline modulations in general skills (Figure 2, upper row)

Considering general skill measurements, the model including only the main effect of Block best fitted both RT and accuracy data, BF_M_ = 5.47 for RT, BF_M_ = 4.64 for accuracy. Main effect of Block was significant and associated with a strong evidence in favor of the effect for RT and accuracy, *F*(1, 114) = 693.32, *p* < .001, η^2^p = .86, BF_10_ = ∞ and *F*(1, 114) = 55.46, *p* < .001, η^2^p = .33, BF_10_ = 1.90 × 10^8^, respectively. RT decrease and accuracy increase over between-block rest periods suggested ultra-fast offline improvements in general skills. Neither main effect of Between-blocks rest duration nor its interaction with Block were significant (for main effects of Block, Fs < 1 for both RT and accuracy measures; for Block × Between-blocks rest duration, *F*(1, 114) = 1.11, *p* = .30 for RT and *F*(1, 114) = 3.28, *p* = .07 for accuracy) and were associated with ambiguous information (for the main effect of Between-blocks rest duration, BF_10_ = 0.45 for RT and BF_10_ = 0.57 for accuracy; for the interaction of Block × Between-blocks rest duration, BF_10_ = 0.45 for RT, BF_10_ = 1.12 for accuracy). These results suggest that ultra-fast offline improvements occurred during between-block rest periods, but no evidence for an effect of between-block rest duration on general skills was observed. In summary, general skills seemed to benefit from short rest periods independently of their durations.

### Ultra-fast offline modulations in probabilistic sequence knowledge (Figure 3, lower row)

For probabilistic sequence knowledge measurements (i.e., the difference between low- and high-probability triplets), the model including only the main effect of Block best fitted both RT and accuracy data (BF_M_ = 11.83 for RT, BF_M_ = 6.77 for accuracy). Main effect of Block was significant and associated with a strong evidence in favor of the effect for RT and accuracy, *F*(1, 114) = 37.12, *p* < .001, η^2^p = .25, BF_10_ = 1.81 × 10^6^ and *F*(1, 114) = 7.85, *p* = .006, η^2^p = .06, BF_10_ = 3.86, respectively. RT indices increased, and accuracy measures decreased over between-block rest periods, suggesting that probabilistic sequence knowledge decayed during between-block rest periods. For both RT and accuracy measures, neither the main effect of Between-blocks rest duration nor its interaction with Block were significant, all Fs < 1, except for the interaction of Block × Between-blocks rest duration for RT, *F*(1, 114) = 2.60, *p* = .11. They were associated with moderate evidence for the null hypothesis (for the main effect of Between-blocks rest duration, BF_10_ = 0.23 for RT and BF_10_ = 0.23 for accuracy; for the interaction of Block × Between-blocks rest duration, BF_10_ = 0.17 for accuracy), except for the interaction of Block × Between-blocks rest duration for RT that was associated to ambiguous information, BF_10_ = 0.52. These results suggest a decay of probabilistic sequence knowledge that does not seem to be influenced by between-block rest period duration.

**Figure 3.**
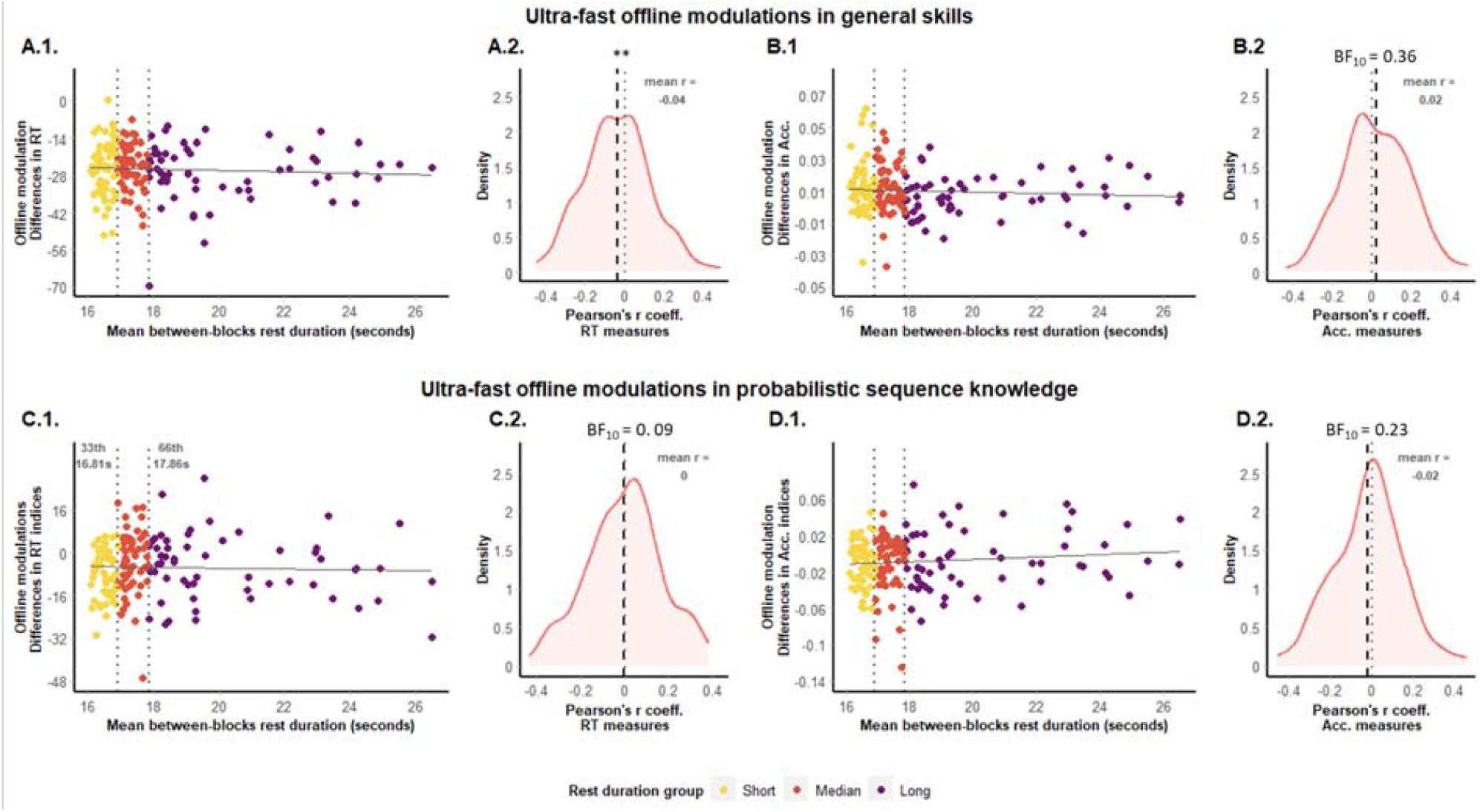
Offline modulation in general skills (top row) and probabilistic sequence knowledge (bottom row) as a function of between-blocks rest duration is reported for response times (A, C) and accuracy (B, D). *(A.1, B.1, C.1, D.1).* Distribution on mean offline modulation depending on mean between-blocks rest duration. Solid black lines represent linear trends and dotted grey lines represent 33^rd^ and 66^th^ percentiles used for the post-hoc subgrouping procedure on mean between-blocks rest duration. Between-blocks rest duration groups resulting from post-hoc subgrouping are color-flagged. *(A.2, B.2, C.2, D.2).* The density of Pearson’s r coefficients resulting from the correlation between between-blocks rest duration and offline modulation for each participant. Dashed lines represent the mean of Pearson’s r coefficients across participants. Dotted lines mark the value to which Pearson’s r coefficients are compared (i.e., zero). These figures highlight the absence of a linear relationship between offline modulation and between-blocks rest duration at the between-participants level for all offline metrics (i.e., general skills, probabilistic sequence knowledge, RT and accuracy measures). However, analyses at the within-participant level show a relationship between offline modulations in general skills and between-block rest periods for both RT and accuracy. Yet, no relationship between offline modulations in probabilistic sequence knowledge and between-block rest duration emerged at the within-participant level. ** stands for p<.01. Bayesian factor are reported for non-significant effects. BF < ⅓ shows evidence for the null hypothesis (see method section for more details).

Taken together, these results suggest that ultra-fast offline improvements occurred in general skills but not in probabilistic knowledge. On the contrary, probabilistic sequence knowledge decayed over the rest period. In both cases, the duration of rest-periods did not seem to influence offline modulations of knowledge. However, the post-hoc subgrouping procedure leads to the categorization of a continuous predictor, which might result in a loss of power (Aiken et al., 1991). Follow up analyses were thus performed to investigate further the relationship between offline decay in probabilistic sequence knowledge and between-block rest periods duration.

### Is there a linear relationship between between-block rest periods and offline modulations in general skills and probabilistic sequence knowledge?

#### Between-participant analysis of offline modulations

For general skills measures, correlations were not significant for RT nor for accuracy and were associated with moderate evidence in favor of the null hypothesis for RTs and accuracy, *r*(170) = −0.06, *p* = .44, BF_10_ = 0.12, and *r*(170) = −0.06, *p* = .40, BF_10_ = 0.14, respectively (Figure 3.A.1 and 3.B.1). For probabilistic sequence knowledge measures, correlations were neither significant for RT nor for accuracy and were associated to strong evidence in favor of the null hypothesis for response times, *r*(170) = −0.03, *p* = .70, BF_10_ = 0.10 (Figure 3.C.1) and moderate evidence for the null hypothesis for accuracy, *r*(170) = 0.10, *p* = .19, BF_10_ = 0.22 (Figure 3.D.1). These results suggest no linear relationship between mean between-blocks rest duration and offline improvement in general skills nor with offline decay in probabilistic sequence knowledge.

#### Within-participant analysis of offline modulations

The absence of a relationship between ultra-fast offline modulations and between-blocks rest duration could be due to high inter-participants variability. To account for the inter-individual variability, we further inspected the strength of the relationship between offline modulation in general skills and probabilistic sequence knowledge and between-blocks rest duration for each participant. For general skill measures, correlation coefficients significantly differed from zero and BF_10_ showed moderate evidence for the alternative hypothesis for RT measure, *t*(171) = - 2.88, *p* = .005, BF_10_ = 4.56 (Figure 3.A.2) and were inconclusive for the accuracy measure, *t*(171) = 1.71, *p* = .09, BF_10_ = 0.36 (Figure 3.B.2). These results suggest a linear relationship between ultra-fast offline improvements in general skills and between-blocks rest duration at the individual level. For the measures of probabilistic sequence knowledge, correlation coefficients did not significantly differ from zero, and BF_10_ showed strong evidence for the null hypothesis for RT measure, *t*(171) = −0.069, *p* = .95, BF_10_ = 0.09 (Figure 3.C.2) and moderate evidence for the null hypothesis for accuracy measure, *t*(171) = −1.42, *p* = .16, BF_10_ = 0.23 (Figure 3.D.2). These results strengthen the lack of a linear relationship between offline decay in probabilistic sequence knowledge and between-blocks rest duration, even at an individual level.

#### Between-blocks rest periods and probabilistic sequence knowledge acquired during the entire experiment

To test whether between-blocks rest duration had a more general influence on probabilistic sequence learning throughout the course of the experiment, we investigated the relationship between between-blocks rest duration and probabilistic sequence learning (i.e., probabilistic sequence knowledge acquired during the entire experiment). Probabilistic sequence learning measures were computed based on mean accuracy and median RT only for correct responses for each participant across the experiment (see Method section).

Beforehand, we ran one-sample frequentists and Bayesian t-tests comparing probabilistic sequence learning to zero to ensure that participants indeed learned probabilistic properties of the sequences during the experiment. Both RT and accuracy scores for probabilistic sequence learning showed significant learning over the experiment and were associated to strong evidence in favor of the alternative hypothesis (for RT: *t*(171) = 24.66, *p* < .001, Cohen’s *d* = 1.88, BF_10_ = 1.352×10^60^, for accuracy: *t*(171) = 19.91, *p* < .001, Cohen’s *d* = 1.52, BF_10_ = 9.40×10^42^).

Then, frequentist Pearson’s correlations and Bayesian correlations between the mean between-blocks rest duration over the task and the probabilistic sequence learning scores were computed. Correlations were neither significant for RT nor for accuracy and associated to strong evidence in favor of the null hypothesis, *r*(170) = 0.03, *p* = .71, BF_10_ = 0.10 for RT and *r*(170) = 0.02, *p* = .77, BF_10_ = 0.10 for accuracy. Similar results were obtained with analyses on probabilistic sequence knowledge acquired at the end of the experiment instead of probabilistic sequence learning averaged over the entire experiment (see Supplementary materials). These results show that while participants have learned the probabilistic structure of the sequences during the experiment, the amount of probabilistic sequence knowledge acquired during the task was not related to between-blocks rest duration.

To sum up, our results showed improvements in general skills but decrements in probabilistic sequence knowledge over between-blocks rest periods. Post-hoc subgrouping variance analyses and between-participants correlational analyses suggested that between-block rest duration was not related to ultra-fast offline improvements in general skills nor to offline decrements in probabilistic sequence knowledge. However, at an individual level, our analyses provided evidence for a positive relationship between rest duration and ultra-fast offline improvements in general skills. Yet, no relationship between rest duration and probabilistic sequence knowledge was observed despite the fact that probabilistic sequence knowledge was acquired during the experiment.

## Discussion

The present study investigated whether short rest periods influence different learning processes. We used an implicit probabilistic sequence learning task that enabled us to distinguish probabilistic sequence learning (i.e., performance depending on probability of occurrence of items) from general skill learning (i.e., performance regardless of the probability of occurrence of items, Hallgató et al., 2013; Nemeth et al., 2010; Song et al., 2007). Participants were allowed to rest after each block of trials, and the beginning of the next block was triggered by them, producing a self-paced fluctuation in the duration of rest periods. The performance was assessed before and after each rest period, granting measures of ultra-fast offline modulation in both general skills and probabilistic sequence knowledge. We wondered (1) whether ultra-fast offline improvements in general skill and probabilistic sequence learning can emerge during between-block rest periods and (2) whether the *duration* of between-block rest period affects offline modulations in general skills and implicit probabilistic sequence knowledge. In other words, can longer rest periods lead to better (or worse) learning performance? We observed that rest periods led to ultra-fast offline increase in general skills and a decrease in probabilistic sequence knowledge. At the group level, neither ultra-fast offline improvements in general skills nor decrements in probabilistic sequence knowledge were linked to rest duration. However, within-participant analyses highlighted that between-block rest duration was related to ultra-fast offline improvements in general skills but not to decrements in probabilistic sequence knowledge.

First of all, our results highlight ultra-fast offline improvements in general skills but a decrement in probabilistic sequence knowledge during rest periods. On the one hand, ultra-fast offline improvements in general skills are in line with a previous study focusing on the learning of deterministic sequences (Bönstrup et al., 2019). On the other hand, ultra-fast offline decrements in probabilistic sequence knowledge seem to oppose a previous study suggesting that memory consolidation of probabilistic information benefit from two-minutes rest periods (Du et al., 2016). Du et al. (2016) showed that offline learning drove the fast acquisition of probabilistic sequences, whereas online learning did not contribute to probabilistic sequence acquisition. This suggested that implicit probabilistic sequence learning couldn’t develop without offline learning. Yet, in our study, probabilistic sequence knowledge was acquired over the experiment despite any evidence for ultra-fast offline improvements, suggesting that implicit probabilistic sequence knowledge *can* develop without it. Several aspects differed between our study and the Du et al.’s (2016): the implementation of probabilistic sequences (a predetermined sequence hidden in random elements in ours, a sequence based on a Markov chain transitional matrix in Du et al.’s), the number of blocks of trials containing to-be-learned probabilistic events (45 blocks in our study, four blocks in Du et al.’s study), the duration of rest periods (self-paced and lasting 18.35 ± 9.40 seconds in ours, fixed at two minutes in Du et al.’s), and the assessment of the probabilistic sequence knowledge (difference between high- and low-probability events in ours, RT measures for the more probable events without comparing them to the least probable). The latter aspect can plausibly explain the discrepant results between Du et al.’s and our study. In Du et al., offline improvements in probabilistic knowledge were not distinguished from improvements in general skills. An offline improvement in general skills (as observed in our study and in Bönstrup et al., 2019) might have influenced the measure of offline learning in probabilistic sequence knowledge. In our ASRT tasks, the measure of probabilistic sequence learning (i.e., difference score between high- and low-probability events) allows to disentangle probabilistic knowledge from general skills (Hallgató et al., 2013; Nemeth et al., 2010; Song et al., 2007). Measures of probabilistic sequence knowledge used in our study thus reflected processes involved in *pure* probabilistic learning, distinct from those underlying general skill learning.

Differences in the duration of rest periods between the two studies might suggest that (1) a crucial parameter might be the duration of rest periods, and (2) a minimum amount of time might be necessary for memory consolidation to take place. Fortunately, our design allowed us to directly test this hypothesis. Beyond the between-blocks rest periods, the experimental design contained two between-session rest periods (mean = 4.30 minutes, SD = 1.66, range = 2.38 – 17.11 minutes) during which participants filled questionnaires. Even if these rest periods were longer, no significant offline learning in probabilistic sequence knowledge emerged (see Supplementary materials). The duration of rest periods does not seem crucial for offline improvements in probabilistic sequence knowledge. To determine in which conditions do offline learning during implicit probabilistic sequence learning emerge and test the hypothesis of a critical period that is essential for offline learning to emerge, future studies should directly manipulate the duration of rest periods, from seconds to a few minutes.

Secondly, we tested whether ultra-fast offline modulations (i.e., improvements or decrements) depends on the duration of between-block rest periods. Concerning ultra-fast improvement in general skill learning, between-participants analyses suggested no influence of the length of between-block rest period on ultra-fast offline modulation neither for general skills. However, within-participants analyses revealed that the longer rest period duration, the stronger ultra-fast offline improvements. In other words, at the individual level, more extended offline periods lead to better learning performance. Our study itself does not allow stating for the mechanisms underlying ultra-fast offline learning of general skills. The index of general skills used in ASRT tasks is a complex measure that encompasses various perceptual and motor activities (e.g., motor-motor and perceptual-motor coordination) and cognitive processes (e.g., adaptation to the task situation; Hallgató et al., 2013; Nemeth et al., 2010; Song et al., 2007). Ultra-fast offline improvements in general skills could be due to the benefit of rest period over either of these processes. Moreover, fatigue or inhibition release can also induce offline improvements (e.g., Brawn et al., 2010; Rickard et al., 2008). By measuring differences in performance, ASRT allows us to rule out the inhibition/fatigue release hypothesis for implicit probabilistic sequence learning measures (Török et al., 2017). Yet, general skills are measured via raw response times and accuracy, thus preventing to eliminate the fatigue effect hypothesis. Ultra-fast offline improvements in motor sequence learning have been linked previously to reactivation of memory traces (Bönstrup et al., 2019; Robertson, 2019). Consistently, one potential explanation of the correlation between rest duration and general skill learning at the individual level in our study is that longer breaks give more time for reactivation or replay neural mechanism to develop during the rest period. Further studies will be necessary to disentangle specific contributions of fatigue release and consolidation processes to ultra-fast offline improvements in general visuomotor skills.

Between-participants and within-participants analyses of the relationship between rest period duration and ultra-fast offline improvements in general skills were inconsistent. From a methodological perspective, this discrepancy seems important for future investigation of the time-dependency of ultra-fast offline improvements. Studies investigating offline processes across longer rest periods operated with a wide range of rest durations (e.g., from one hour to half a day, Press et al., 2005). On the contrary, investigating the time-dependency of ultra-fast offline consolidation does not allow such variability in rest period durations. For example, in the present study, standard (i.e., not outlying) between-block duration ranged from 15.39 seconds to about two minutes. This might have led to a lack of sensitivity of duration measures at the group level. As highlighted in the present study, within-participant analyzing methods can uncover effects due to time that failed to be measured by between-participant analyses. We thus recommend that future studies investigate rest period duration effect on ultra-fast offline improvements at participant level.

Concerning probabilistic sequence measures, we observed a decrease of knowledge over rest period duration and, consistently at the between-participants and within-participants level, these ultra-fast offline decrements were not related to the between-block rest duration. In other words, longer averaged offline periods did not lead to stronger forgetting. This result raises the question of what causes forgetting in probabilistic knowledge during short rest periods. Forgetting can be due to two processes: time-based decay and interference. Decay theory posits that memory traces fade away with the mere passage of time (Brown, 1958), but this theory is still widely debated (Ricker et al., 2014). Other studies suggest that in implicit probabilistic learning studies, interference contributes to forgetting to a great extent because events are typically generated by recombining a small number of features, thus strongly interfering with each other (Perruchet & Pacton, 2006). In our study, ultra-fast offline decrements in probabilistic sequence knowledge were not related to the duration of the rest period, suggesting that probabilistic sequence knowledge did not decay during short rest periods. In ASRT tasks, random events are based on the same features as pattern elements (i.e., spatial location and its mapping to the response keys) and are likely to interfere with pattern elements. Forgetting over the rest periods observed in the present study thus seems more likely to come from interference, which might explain the absence of time-based decay. Future studies will need to disentangle the contribution of time and interference in forgetting during implicit probabilistic learning. To do so, we suggest to orthogonally manipulate the rest period duration and the amount of interference between target and not-target event.

Consistently with Bönstrup et al. (2019), general skills improve over a short period of time. Further, longer duration of rest periods leads to stronger improvements, suggesting that more consolidation processes, such as memory replay, might have taken place. Implicit probabilistic sequence knowledge, however, is prone to forgetting rather than offline learning over short periods within a single training session. Forgetting of probabilistic sequence knowledge does not seem to depend on time, and might rather be due to interference. The opposing results for general skills and implicit probabilistic learning suggest differences in mechanisms underlying memory consolidation of deterministic and probabilistic information. Because of the shortness of rest periods, our results raise the question of a critical time period for consolidation to occur and compensate or overcome forgetting of implicit probabilistic knowledge. Future studies investigating the time-dependency of consolidation processes underlying ultra-fast improvements should manipulate the duration of rest periods and account for within-participant variability. Beyond the question of a crucial time period, we question the mere existence of ultra-fast consolidation processes in probabilistic learning. Future studies should carefully distinguish processes underlying general visuomotor learning, task adaptation, and fatigue/inhibition release from those involved in implicit probabilistic learning.

## Acknowledgements

This research was supported by the IDEXLYON Fellowship of the University of Lyonas part of the Programme Investissements d’Avenir (ANR-16-IDEX-0005) (to D.N.); National Brain Research Program (project 2017-1.2.1-NKP-2017-00002); Hungarian Scientific Research Fund (NKFIH-OTKA K 128016, to D.N., NKFIH-OTKA PD 124148 to K.J.); János Bolyai Research Scholarship of the Hungarian Academy of Sciences (to K.J.). The authors are thankful to Kate Schipper for proofreading the manuscript.

## Contributions

DN, LF, RQ conceived the presented idea, KJ and DN designed the study and organized the data collection. LF and CP analyzed the data. LF wrote the first draft of the manuscript. TV, RQ, KJ, and DN reviewed and critically edited the previous versions of the manuscript. All authors read and approved the final version of the manuscript.

