## Supplementary Material for "The longer the better? General skill but not probabilistic learning improves with the duration of short rest periods" (from another version of this preprint: version 1, 2020-05-14)

**Between-group differences in demographic variables**

The between-group differences (Short rest duration group vs. Long rest duration group) in demographic variables (Age, Education level, and Sex) were assessed with frequentists and Bayesian one-way ANOVAs for continuous measures and chi-squared contingency tables test for categorical measures.

Table 1. Overview of demographic characteristics for the two participants’ groups resulting from the post-hoc subgrouping procedure. Descriptive statistics are mean (standard deviation) for continuous measures and frequency distribution for categorical measure. Between-groups comparison refers to the result of one-way ANOVAs and Chi-Squared contingency table comparing between-group performance for continuous and categorical variables, respectively. No between-group differences in Age, Education level or Sex emerged.

| **Measures** | **Descriptive statistics** | | **Between-groups comparison** |
| --- | --- | --- | --- |
|  | Short rest period group – N = 57 | Long rest period group – N = 59 |  |
| Rest duration | 16.52 (0.22) seconds | 20.34 (2.25) seconds | N. A. |
| Age | 21.60 (4.12) years-of-age | 21.72 (4.26) years-of-age | F<1, BF = 0.20 |
| Education level | 14.54 (2.39) years | 14.78 (2.00) years | F<1, BF = 0.23 |
| Sex | 47 females and 10 males | 51 females and 8 males | χ² = 0.35, p = .55, BF = 0.20 |

**Analyses of statistical knowledge acquired at the end of the experiment**

Probabilistic sequence learning acquired by the end of the experiment was measured on the mean accuracy and median RT only for correct responses during the last five blocks for each participant (for more details on the measures computation, see the Methods section), further referred to as final probabilistic sequence knowledge.

To ensure that participants had significant probabilistic sequence knowledge at the end of the experiment, we ran one-sample frequentists and Bayesian t-tests comparing probabilistic sequence knowledge to zero. RT and accuracy measures of final probabilistic sequence knowledge showed significant knowledge and were associated to strong evidence in favor of the alternative hypothesis (for RT: *t*(171) = 19.80, *p* < .001, Cohen’s *d* = 1.51, BF_10_  = 4.88×10^42^; for accuracy: *t*(171) = 12.85, *p* < .001, Cohen’s *d* = 0.98, BF_10_  = 5.28×10^23^).

Then, frequentist and bayesian Pearson’s correlations between the mean between-blocks rest duration over the task and the final probabilistic sequence knowledge measures were computed. Correlations were not significant for RT nor for accuracy and were associated to weak evidence for the null hypothesis for RT, *r*(170) = 0.06, *p* = .46, BF_10_  = 0.12, and strong evidence for null hypothesis for accuracy, *r*(170) = 0.02, *p* = .83, BF_10_  = 0.10. These results are in line with analyses on probabilistic sequence knowledge averaged over the experiment and comfort that while participants seem to have learned the probabilistic structure of the stimuli during the experiment, the amount of probabilistic learning was not related to between-blocks rest period duration.

**Analyses of offline learning during between-sessions rest periods**

To test whether implicit probabilistic sequence knowledge improved during longer rest periods, we analyzed offline improvements during between-sessions rest periods (two periods per participant, mean = 4.30 minutes, SD = 1.66, range = 2.38 – 17.11 minutes). Measures of offline modulations in probabilistic sequence knowledge during between-sessions rest periods followed the same procedure as for between-blocks rest periods (see Method section for a more detailed description). First, we measured mean accuracy and median RT for correct responses for high- and low-probability triplets of the first two sequences and the last two sequences of each block. This resulted in four RT scores and four accuracy scores for each block (high-probability triplets and low probability triplets in the two first sequences and in the two last sequences). Then, we computed accuracy and RT indices of probabilistic sequence knowledge (i.e., score for high-probability triplets minus score for low probability triplets) for the two first sequences and the two last sequences of each block. Finally, the measure of offline modulations consisted of the difference between the index of probabilistic sequence knowledge of the first two sequences of a given block minus the index of probabilistic sequence knowledge of the last two sequences of the previous one. This procedure resulted in two scores (for RT and for accuracy) for each block (i.e., two per participant), that were averaged for each participant. For RT indices, a negative score shows a speeding up of responses and suggests offline improvement, while a positive score shows a slowing down of responses and suggests memory decay. For accuracy, the opposite pattern is true.

We ran one-sample frequentists and Bayesian t-test comparing scores of offline modulations in probabilistic sequence knowledge to zero, to test whether probabilistic sequence knowledge either improved or decreased during between-sessions rest periods. Neither RT nor accuracy scores for probabilistic sequence learning showed significant offline modulations in during between-sessions rest periods (for RT: t(173) = 1.96, p = .052, Cohen’s *d* = .15; for accuracy: t(173) = -0.98, p = .33, Cohen’s *d* = -.07) and were associated to ambiguous evidence for RT, BF_10_ = 0.54, and moderate evidence for the null hypothesis for accuracy, BF_10_ = 0.14. These results suggest no offline improvement nor offline decrement in implicit probabilistic sequence knowledge during between-sessions rest periods.
